# Depth-Dependent Alterations in Cerebral Blood and Tissue Oxygenation Associated with Tau in Alzheimer’s Disease Models

**DOI:** 10.64898/2026.10.09.757981

**Authors:** Nancy E. Ruiz-Uribe, Mohammed Alfadhel, Nina Wolf, Sophia Esperanza Murray, Elaina Gross, Ava Devine, Danielle G Shaby, Simin Belali, Sergei Vinogradov, Theodore Zwang, Martin Thunemann, Sava Sakadzic, Rachel E Bennett

## Abstract

Vascular dysfunction is increasingly recognized as a feature of Alzheimer’s disease (AD), and inadequate oxygen delivery may contribute to neuronal death. However, the relationship between cerebral oxygenation and neuronal pathology remains unclear. Here, we used two-photon phosphorescence lifetime microscopy (2PLM) with Oxyphor2P to measure intravascular and tissue partial pressure of oxygen (pO2) at three cortical depths (0, 100 and 200 µm) in awake rTg4510 and THY-Tau22 mice. In 9-month-old rTg4510 mice, capillary and venular pO2 were lower than in WT, mainly in deeper layers, and oxygen extraction fraction was increased, whereas 6-month-old mice showed increases in capillary and venular pO2. Suppressing the tau transgene with doxycycline for three months did not restore vascular pO2. THY-Tau22 mice, despite limited tangle pathology, showed even greater reductions in capillary pO2 at every depth. In 6-month-old rTg4510 mice, tissue pO2 trended lower than in WT mice, and the fraction of low pO2 values (< 18 mmHg) increased with depth, reaching 33% at 200 µm versus 4% in WT. By labeling neurofibrillary tangles (NFTs) with the tau-binding dye HS84, we found that tissue pO2 was significantly lower in tangle-positive areas at 100 and 200 µm. Longitudinal imaging of 2,282 neurons showed that disappearing neurons were not located in relatively hypoxic tissue within an animal. Finally, ex vivo immunostaining revealed neuronal loss and increased astrogliosis in rTg4510 mice, while vascular architecture was unchanged. Altogether, our results indicate that tau disrupts cerebrovascular and tissue oxygenation in a depth- and age-dependent manner, and that neuronal loss is not explained by the oxygen levels of a neuron’s immediate microenvironment but may instead reflect a global reduction in cortical oxygenation.

## INTRODUCTION

Vascular dysfunction is increasingly recognized as a feature of Alzheimer’s disease (AD)^1–3^. Human neuroimaging studies, for example, show that cerebral perfusion is reduced in individuals with AD even at early stages^4^. Since the brain is a highly active metabolic organ that requires uninterrupted blood flow to maintain neuronal activity^5^, changes in cerebral blood flow will alter oxygen supply and can have a profound effect on neurodegeneration^6,7^. Consistent with this, in addition to reduced CBF, both oxygen metabolism^8^ and neurovascular coupling^9,10^ are also altered in AD. However, it is not clear if these vascular changes directly contribute to the loss of neurons in disease or are the result of reduced metabolic demands that arise with the progression of disease.

Technological limitations, specifically, the lack of techniques to measure oxygenation with high spatial and temporal resolution, have prevented the study of *in vivo* oxygen dynamics in both human brain and in AD mouse models. While recent neuroimaging studies in humans shows that cerebral blood flow is closely associated with the accumulation of pathological tau protein in the brain and cortical atrophy, these studies lack the ability to directly visualize cells and cannot directly quantify how differences in oxygen distributions throughout the tissue are impacted by the presence of neurons with tau aggregates and/or whether changes in oxygenation influence cell loss. The goal of this work is to test the hypothesis that Alzheimer’s-related neuronal death is, in part, driven by inadequate vascular support and oxygenation.

To test this hypothesis, we used two-photon phosphorescence lifetime microscopy (2PLM)^11^ to directly measure oxygen delivery^12–14^ both within individual vessels and throughout the cortex of awake mice. Partial pressure of oxygen (pO2) was measured with Oxyphor2P^15^, a phosphorescent probe with strong two-photon absorption near 960 nm and high emission quantum yield that enables 3D longitudinal pO2 measurements. We examined both the rTg4510 mouse line, which develops robust tangle pathology throughout the cortex at young ages, and the THY-Tau22 line that has elevated tau production by limited tangle formation. By co-labelling the vasculature with FITC-dextran and neurofibrillary tangles (NFTs) with HS84^16^, we can establish relationships between vascular and tissue pO2 to NFTs. Here, we provide new evidence that the accumulation of tau pathology in the brain disrupts normal cerebrovascular supply and contributes to lowering of pO2 values (pO2 < 18 mmHg^17^) in cortical areas. In addition, we show that tau tangles and neuroinflammation driven by astrogliosis, rather than neurons, are associated with altered oxygen consumption observed in the mouse cortex at depths between 0 and 200 µm.

## METHODS

### Mice

All animal procedures were performed in accordance with the protocols of the Institutional Animal Care and Use Committee (IACUC) at Massachusetts General Hospital. An initial cohort of 6-month-old (n = 6) and 9-month-old (n = 6) rTg4510 mice, along with their corresponding WT controls (n = 4), was used for vascular pO2 measurements and immunostaining. A second separate cohort of 6-month-old rTg4510 and WT mice (n = 4 per group) was used for tissue pO2 measurements and immunostaining. A third separate cohort of 6-month-old rTg4510 (n = 4) and WTs (n = 4) was used for the neuron loss experiments. The rTg4510^18^ mice were obtained from Jackson Laboratories (Cat # 024854). We used 11-month-old THY-Tau22 with their corresponding WT controls (n = 4 per group) for vascular pO2 measurements. A separate cohort of THY-Tau22 mice (11 months, n = 6) was used for immunostaining. The THY-Tau22 mouse line^19^ was originally obtained from Luc Buee and is maintained in-house via breeding THY-Tau22 male mice to C57BL/6J females from Jackson Laboratories (Cat # 000664). The two initial cohorts of rTg4510 and THY-Tau22 mice with their corresponding WT littermates were used for blood volume measurements. Only rTg4510 (no WT) and THY-Tau22 (with WT) mice from those initial cohorts were used for hematocrit measurements. Both female and male mice were used in all experiments. Mice were housed under a 12-hour light/dark cycle and provided food (Lab Diet cat no. RMH3000 5P75) and water ad libitum. For doxycycline treatment, mice were provided with a diet containing 200 mg/kg doxycycline (Bio-Serv cat no. S3888) for three months.

### Cranial window surgeries

The implantation of cranial windows was done as follows: First, mice were anesthetized with 100 mg/kg of ketamine and 10 mg/kg of xylazine or isoflurane (4% for induction, 1.5% for maintenance), body temperature was maintained at 37°C with a heating pad. Lidocaine was injected into the scalp, and then an incision was made along the midline. The skin was glued around the scalp with tissue glue, and a craniotomy was performed over the somatosensory cortex of 5 mm in diameter. A sterile 5 mm glass coverslip with a silicon port^20^ was placed over the craniotomy and then glued with metabond over the skull. A metal headpost was then glued with metabond at the site of the craniotomy. Mice were given a 3.25 mg/kg body weight dose of Ethiqa XR (Fidelis) and Tylenol in water (300 mg/ 100 ml) and allowed to recover on a heating pad until conscious. Mice were ready to start habituation and imaging after three weeks post-surgery.

For the labelling of neurons with mKalama, we generated a plasmid to ligate the mKalama1-Nuc sequence to an AAV backbone containing the human synapsin1 promoter, and mass produced it at a final titer concentration of 1.53E+13 GC/mL as described in detail in Zwang et al.^21^ During cranial window surgery, before implanting the glass windows, mice were injected with 1 µL of AAV-mKalama using a syringe pump (Harvard Apparatus) equipped with a Hamilton syringe at a speed of 0.2 µL/min, targeting 0.3 mm below the surface of the cortex, and the needle was left in place 5 min post-injection to ensure correct diffusion of the viral vector. The brain was kept moist with gel foam soaked in sterile 1X PBS during the entire procedure. After the injection, the mice were implanted with a glass window containing a silicon port and a metal head post to facilitate awake imaging. Animals were then allowed to recover for at least three weeks to reduce neuroinflammation and allow full expression of the fluorophore.

### Habituation to awake imaging

After mice were recovered from surgery, they were habituated to head fixation daily for a period of 4 days, starting with a 15-minute session and in gradual increments of 15 minutes until reaching one hour of total head restraint under the microscope objective. Mice were briefly placed in a chamber containing 1.5% isoflurane, and once asleep, they were head-restrained on a stereotax with a rotating wheel where their bodies were allowed to move freely. After the end of the habituation period, a small amount of food reward was given to the mice.

### Delivery of optical probes

Two separate protocols were used depending on the type of measurement performed (vascular or tissue pO2). For vascular measurements, mice were retro-orbitally injected with a 50 µL solution of Oxyphor2P at a final concentration of 5 µM, which was diluted with sterile 1X PBS from a 250 µM stock solution based on the animal weight as described in detail by Erlebach et al^14^. For tissue (extravascular) measurements, Oxyphor2P at a concentration of 250 µM was delivered through the silicon port via intracortical microinjection (Nanoject II) with a glass pipette. Pipettes were pulled using a P-200 Sutter Instrument puller (O.D 1.00 mm, Sutter Instrument), filled with the Oxyphor2P solution and fixed using a micromanipulator at an angle of approximately 45 degrees. Mice were anesthetized with isoflurane and placed in a stereotaxic frame. The tip of the pipette was visualized through a stereotaxic microscope and guided towards the port surface. The pipette was then introduced through the port for approximately 300 µm deep into the cortical surface. Two injections of 50 nL were performed at 200 µm and 100 µm. We waited 5 minutes in between injections to allow the dye to fully diffuse into the brain tissue. After the injections, mice were retro-orbitally injected with 50 µL of FITC-Dextran (70kDa) and allowed to recover for approximately ∼1h until the start of the imaging session. For labeling neurofibrillary tangles with HS84 (a blood-brain-barrier-penetrant luminescent conjugated oligothiophene dye)^16^, mice were retro-orbitally injected with 150 µL of a 150µM solution of HS84 reconstituted in 1X PBS, three to four days before imaging, to allow diffusion of the dye and clearance of unbound dye from the vasculature.

### Two-photon phosphorescence lifetime microscopy (2PLM)

Imaging was performed using a Ultima2P two-photon laser scanning microscopy system (Bruker) with a Ti:Saphire laser (Insight X3, SpectraPhysics). An Olympus 20X objective (UMPlanFI, NA = 1.05) was used for data collection and imaging, and an objective heater (Okolab) was used to warm the objective to 45°C. A thermocouple (Fisher Scientific) was used to measure the temperature of the ultrasound gel between the objective and the cranial window, ensuring it remained constant at 36°C-37°C, so that correct calibration parameters could be used to covert phosphorescence lifetimes of Oxyphor2P to pO2 values^15^. As in a previous study by Machler et al., the emission of Oxyphor2P was reflected by a low-pass dichroic mirror with a 900-nm cutoff (Chroma), then pass through a 795/150-nm bandpass filter (Semrock) and detected using a photon-counting photomultiplier tube (PMT; H10770A-50, Hamamatsu). A second PMT (H10770B-40, Hamamatsu) with a 525/50 nm bandpass filter (Semrock) was used to detect fluorescence of FITC-labeled dextran and the neurofibrillary tangle dye HS84. Photon counts were detected using a Time-Correlated Single Photon Counting Card (Becker and Hickel,SPC-150).

First, a 3D z-stack was acquired from the top of the cortex to 300 µm to get a vascular map. A 300 µm x 300 µm FOV was selected at each depth (0, 100, 200 µm) for vascular and tissue pO2 measurements. The initial depth of z = 0 µm lies right below the dura mater. Vessels were manually classified as arterioles, venules or capillaries by tracing their connection to pial arterioles and venules, assessing their morphology, vessel diameter, and direction of flow as previously described^23^. For vascular measurements, individual point scans of vessels at each imaging plane were manually selected. Phosphorescence was excited using a 10 µs-long train of femtosecond pulses (960 nm, 110 fs) followed by a collection period of 290 µs. A total of 2000 cycles per point were collected. Measurement points not located within a clearly resolved vessel lumen (e.g., diffuse signal without a defined vessel boundary) were excluded. For tissue measurements, we followed a protocol described by Machler et al.^22^, in which a square grid of 400 points was acquired using the same excitation and collection periods, with 50 cycles per point and 20 repetitions (1000 cycles per point), for a total acquisition time of 120 seconds^22^. Animal motion was carefully monitored using CrownTracker software (MaJn Lab), which records the animal’s position and velocity by tracking wheel movement, and any data in which motion was detected was immediately discarded. Photon counts were binned into 2 µs long bins, and data were analyzed using a custom MATLAB code.

### Image and data analysis for estimation of pO2

Phosphorescence decays were analyzed as described in Erlebach et al^14^. In brief, decays were fitted starting 5 µs after the end of the excitation period using the equation below with a custom MATLAB code:

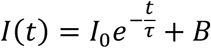

Where *I*(*t*) is the phosphorescence intensity as a function of time, *I*_!_ is the initial phosphorescence signal, t is the probe decay time and B is a constant baseline term. To obtain pO2 values, an empirical equation was used:

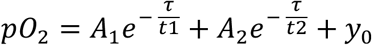

where A1, A2 and y0 are constants obtained from independent calibrations^15^.

### Calculation of oxygen extraction fraction

Hemoglobin oxygen saturation (sO2) was calculated using the Hill equation^24^:

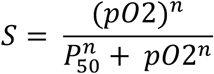

Where P50 corresponds to the partial pressure of oxygen at which hemoglobin is 50% saturated (for mice, P50 = 40.2 mmHg), *n* is the Hill coefficient for C57BL/6 mice (n = 2.59). Oxygen extraction fraction (OEF) was then computed by calculating the average arteriole and venular pO2 per mice per depth, and using the equation^25^:

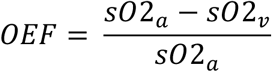

Where sO2a and sO2v correspond to sO2 in arterioles and venules, respectively.

### Correlation of vascular pO2 with perivascular tangle density

Vascular pO2 was acquired using Oxyphor2P (red channel), and tangles were imaged with HS84^16^ (green channel). A two-channel composite image was then generated for each 2D field of view at z = 0, 100, 200 µm. For every vessel in which pO2 had been measured, a circular region of interest of 20 µm radius was drawn manually at the center of the vessel, and the mean integrated density of the HS84 channel (tangles) within that region was measured in ImageJ (v.2.16.0). Each vessel was thereby assigned a single HS84 raw integrated density value, which was plotted against its intravascular pO2 value and correlated in GraphPad Prism (v.11.0.0) using Pearson correlation, separately by vessel type, depth and age.

### Classification of tangle-positive/tangle-negative tissue and correlation with tissue pO2

For tissue pO2, HS84 images were background-subtracted, thresholded (Otsu method) and binarized in ImageJ. The corresponding tissue pO2 grid was then overlaid on the binarized tangle image, and a custom MATLAB script (v.R2026a, MathWorks) assigned each grid point to a tangle-positive or tangle-negative area by majority neighborhood classification within a circular mask of 13-pixel (∼4 µm) radius centered on that point. A point was classified as tangle-positive when at least 3% of the pixels within its mask were HS84-positive, a threshold chosen because it best matched the visible tangle distribution in the scanned area. pO2 values were then averaged across tangle-positive and across tangle-negative points, separately for each depth (z = 0, 100, 200 µm) and animal.

### Cell death tracking with mKalama-labeled neurons

To track neuronal death based on initial tissue pO2 values, a 1X stack was acquired at week 1 in the region where neurons could be visualized. Then, a 2X zoom 50 µm substack was acquired to visualize a subset of the neuronal cluster. A 100-point pO2 grid was then acquired at the top of the neuronal subcluster. Then, the same neuronal subcluster was imaged at week 2, 3 and 4. Neuronal subclusters were then aligned using IMARIS as described in detail in Zwang et al., 2024, and color-coded based on the week of imaging. Only neurons that showed up in week 1 were tracked across multiple weeks, to account for technical limitations that can stem from an uneven field of view or different imaging sessions. Disappearing neurons were manually identified based on alignment and color coding. Tissue pO2 values were assigned to each neuron by using a nearest neighbor algorithm on MATLAB.

### Hematocrit and blood volume measurements

Mice were briefly anesthetized with isoflurane (3%), placed in a sterile field and scruffed. A 16G needle was used to pinch the submandibular vein, and a drop of blood was collected with a glass capillary tube. The tube was sealed with wax on one end and placed on a hematocrit tabletop centrifuge for 15 seconds. Hematocrit (%) was measured by aligning the line where the plasma starts on the tube to the hematocrit chart percentages.

Total blood volume was estimated by using the equation: *BV* = 0.06 *X BW* + 0.77, where BV is blood volume in mL and BW is body weight in grams^26^.

### Immunohistochemistry

Upon reaching the experimental endpoint, mice were euthanized with CO_2_ and transcardially perfused with 1X phosphate buffered saline (PBS). Brains were extracted, cut in half along the midline, stored for 24h in 4% PFA at 4°C and then transferred to 30% sucrose for immunohistochemistry Hemi-brains sliced on a microtome with a 40 µm thickness. Slices were placed on 30% glycerol solution and stored at -20C until further processing.

For immunostaining, free floating sections were washed 3X with Tris-buffered saline (TBS), incubated in blocking solution (3% normal goat serum in TBS with 0.25% Triton-X) for 1 hour, and then incubated in primary antibody overnight at 4C. On the next day, sections were washed 3X with TBS and incubated with secondary antibody for 2h. Sections were then washed one last time 3X with TBS, mounted on glass slides and sealed with FluoromountG + DAPI. Images were acquired with a slide-scanning microscope (Olympus VS120) with a 20x objective and analyzed using QuPath’s positive cell detection algorithm. Layer-by-layer analysis was done by manually segmenting the cortex into 50-µm strips by a blinded experimenter. Antibodies used are as follows: anti-NeuN Alexa Fluor 568, (Cat # ab207282, Abcam), anti-GLUT-1 Alexa Fluor 488, (Cat # 07-1401-AF488, Millipore), anti-AT8 (Cat # MN1020, Invitrogen), Rabbit anti-IBA1 (Cat # 019-19741, FujiFilm Wako Chemicals) conjugated next day with anti-Rabbit Alexa Fluor 564 (Cat # A32740, Thermofisher), and anti-GFAP Alexa Fluor 488, (Cat # 53-9892-82, Thermofisher). All antibodies were used in a 1:500 dilution.

### Measurement of vessel diameter, tortuosity, density and branch length from immunohistochemistry images

GLUT-1 images of blood vessels were cropped to include only the cortical region and were analyzed using ilastik (v.1.4.2)^27^. In brief, object classification of vessels and background was performed manually to train the machine-learning based classifier. Metrics such as tortuosity, mean branch length and vessel density were extracted from ilastik after the segmentation was complete. For diameter measurements, images of vessel segmentations from ilastik were then converted to binary masks on ImageJ, and BoneJ^28^ was used to calculate a distance map from the skeletonized image, providing average vessel diameters that were then converted to µm using a pixel to µm conversion factor.

### Statistics

Data was analyzed with RStudio (v.2026.09.0) and GraphPad Prism. To analyze statistical differences between groups, data was first tested for normality using a D’Agostino Pearsons test. If data was normal between two different groups, a two-tailed t-test was performed, otherwise we used a Mann-Whitney test. We used two-way ANOVA and linear mixed effect models to assess the effect of depth, vessel type and age in vascular pO2 measurements where multiple depth-dependent measurements per animal were performed. To evaluate differences between experimental groups, post hoc multiple-comparison tests were also performed (Tukey HSD or Sidak).

For analysis of Figure 5: tissue pO₂ was measured at 2,282 individual neurons across seven animals (four rTg4510, 290 neurons; three wild-type, 1,992 neurons). The animal was treated as the experimental unit throughout, and all descriptive statistics are animal-weighted (means of per-animal means). Because baseline oxygenation varied several-fold between animals (per-animal means 16.0–46.7 mmHg), pO₂ was expressed as each neuron’s deviation from its own animal’s mean (ΔpO₂, within-animal centering), so that the analysis tests whether disappearing neurons occupy relatively hypoxic tissue within an animal, independent of between-animal differences in baseline oxygenation, imaging session, and cortical depth. The primary model was a mixed-effects logistic regression of disappearance on ΔpO₂, genotype, and their interaction, with a random intercept for animal (lme4 v1.1-35.1, Laplace approximation, BOBYQA optimizer); odds ratios are reported per 10 mmHg of ΔpO₂ and fixed effects were confirmed stable across alternative optimizers. The random-intercept variance was estimated at the boundary once genotype was included, indicating that genotype accounted for the between-animal differences in disappearance rate; the model was therefore also fitted conditionally, with animal as a fixed effect and Firth’s penalized likelihood to correct small-sample bias and guard against quasi-separation (brglm2 v0.9.2, profile-likelihood intervals). Two further estimators were fitted: generalized estimating equations with an exchangeable working correlation and animal as the clustering unit (geepack v1.3.10), interpreted as a directional cross-check only given that robust standard errors are anti-conservative with seven clusters; and a distribution-free permutation test in which disappearance labels were shuffled within each animal (10,000 permutations), using the animal-weighted mean ΔpO₂ of disappearing neurons as the test statistic. Sensitivity analyses re-expressed pO₂ in within-animal standard deviation units and refitted the model on absolute pO₂. Only effects consistent across all four estimators were interpreted. Figures were produced with ggplot2 v3.4.4; all analysis code is available at https://github.com/nancyruizu/po2-neuron-death. P- values less than 0.05 were considered statistically significant and shown in graphs with a “*”, while for values between 0.05 and 0.1 we reported a “trend” in the data.

## RESULTS

### Reductions in capillary and venular pO2 are seen with cortical depth in 9-month-old rTg4510 mice but not in 6-month-old rTg4510 mice

To evaluate intravascular pO2, we used two-photon microscopy to examine the somatosensory cortex of awake wild type (WT) and rTg4510 mice, at 6- and 9-months of age (**Fig. 1A**). These two ages were chosen to examine changes at different stages of disease progression. In this mouse model, NFTs start to form in the brain at 4 months and are accompanied by neuron loss by 6 months. Frequent tangle pathology can be seen from 9 months onwards^29^. First, we obtained a 3D vascular map (**Fig. 1B**) to locate the area of imaging. Starting just below the dura mater (z = 0 µm), we selected 9-10 points of measurements in vessels, including arterioles, venules and capillaries, and took additional measures at z = 100- and 200-micron depths (**Fig. 1C**), given that microvascular densities vary significantly with depth in the mouse cortex^30^ and prior work shows depth-dependent changes in oxygenation^22,31,32^. Individual point scans were performed in each vessel, and phosphorescence decays from Oxyphor2P were subsequently converted into pO2 values (mmHg) using the Stern-Volmer equation (**Fig. 1C, D, E**). This resulted in a total of 464 measured vessels, broken down by age and genotype as follows: In 6-month-old mice: arterioles: n = 46 (rTg4510), n = 24 (WT), venules: n = 19 (rTg4510), n =16 (WT), capillaries: n = 106 (rTg4510), n = 53 (WT). In 9-month-old mice: arterioles: n = 39 (rTg4510), n = 27 (WT), venules: n = 27 (rTg4510), n = 19 (WT), capillaries: n = 58 (rTg4510), n = 30 (WT). In 11-month-old THY-Tau 22 mice, arterioles: n = 41 (WT), n = 25 (THY-Tau22), venules: n = 18 (WT), n = 46 (THY-Tau22), capillaries: n = 62 (WT), n = 97 (THY-Tau22).

**Figure 1.**
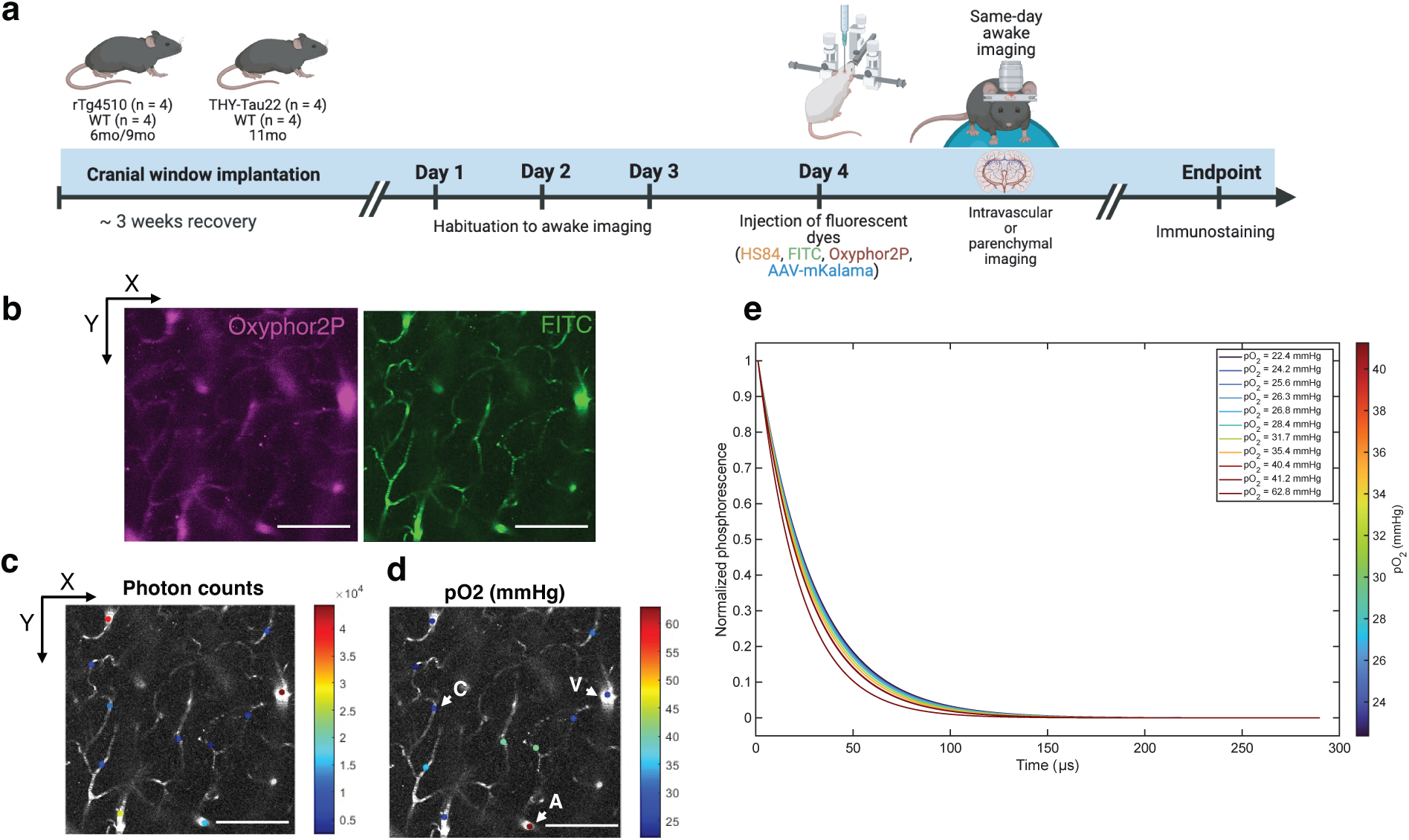
Overall experimental design and methods. **a)** rTg4510 mice (ages 6-mo and 9-mo) and THY-Tau22 mice (aged 11mo) and their corresponding WT littermates were used for awake imaging and pO2 measurements. Mice were implanted with cranial windows, trained for awake imaging, and injected (R.O) with FITC-dextran to label blood vessels, HS84 to label tangles, and Oxyphor2P for intravascular pO2 measurements. For extravascular pO2 measurements, Oxyphor2P was injected I.C through a silicon port. For neuron tracking experiments, AAV-mKalama was injected I.C before cranial window implantation. After imaging was completed, mice were euthanized, and their brains were extracted for immunohistochemistry. **b)** Two-dimensional Z-stack of the brain vasculature in a mouse labeled with FITC-dextran and Oxyphor2P. Scale bar = 300 µm. **c)** Phosphorescent photon counts detected after laser excitation from the different vessels labeled with Oxyphor2P. **d)** pO2 values from vessels in Fig. 1C after converting photon counts using an empirical equation. Capillaries (C), venules (V), and arterioles (A) are labeled. **e)** Normalized individual phosphorescence decays detected during the collection period (290 µs) from each vessel are measured in Fig. 1D with their corresponding pO2 values.

During imaging, an initial 300-µm-deep microvascular angiogram was acquired to obtain an overall vascular map for vessel identification **(Fig 2A).** No significant physiological differences in hematocrit and blood volume were found comparing rTg4510 and WT mice **(Sup Fig 6A,C).** In 6-month-old mice, we assessed vascular pO2 at z= 0, 100 and 200 µm and found a significant increase in capillary pO2 at 200 µm and venular pO2 at 100 µm and 200 µm (**Fig. 2C**). In 9-month-old mice rTg4510 mice, we found significant decreases in capillary pO2 at 0 µm and 200 µm, and venular pO2 at 100 µm and 200 µm **(Fig 2E).** We did not observe significant differences in vascular pO2 heterogeneity as seen by the standard deviation of the mean pO2 values per mice in both rTg4510 and THY-Tau22 mice **(Sup Fig 2, 3).** Since vascular segments along the arterio-venular axis were assessed, we were also able to calculate the average oxygen extraction fraction (OEF) within our imaged cortical region. We found trends towards reduced OEF in 6-month-old rTg4510 mice and a significantly higher OEF in 9-month-old rTg4510 mice at 100 µm deep **(Fig 2F, 2G)**. A lower OEF may indicate increased blood flow while tissue oxygen metabolism remains unchanged. On the other hand, higher OEF may indicate reduced capillary blood flow or increased metabolic demand by the tissue.

**Figure 2.**
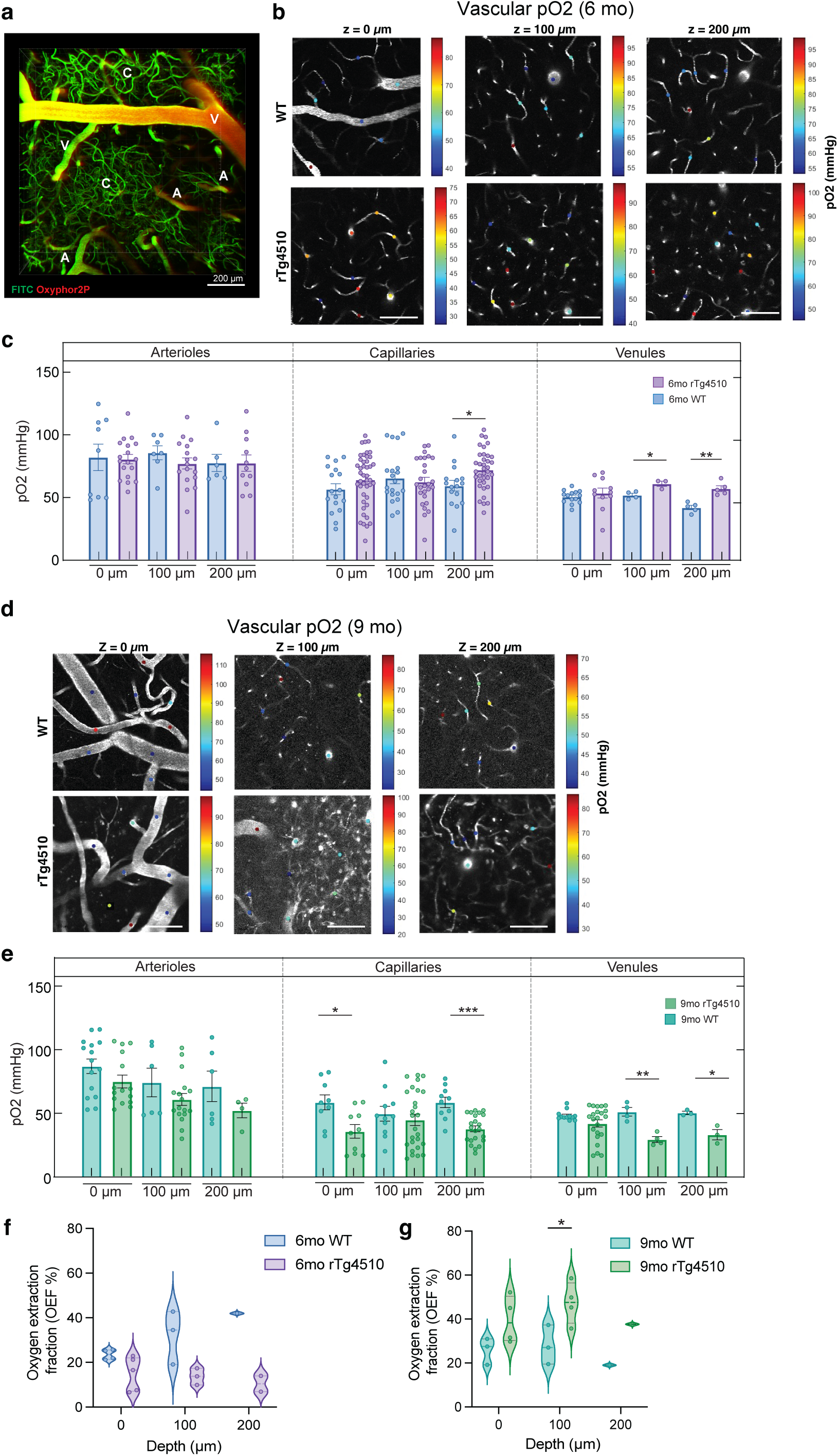
Intravascular measures show increased pO2 in 6-month-old mice and reduced pO2 in 9-month-old rTg4510 mice. **a)** Z-projection of a 3D stack of cerebral vessels labeled with FITC-dextran and Oxyphor 2P. Examples of venules, capillaries, and arterioles are highlighted. **b)** 2D plane images at each depth (0, 100, 200 µm) of vascular segments imaged with 2PLIM where point scans were used to individually measure pO2 values in different vessel types from 6-month-old WT and rTg4510 mice. **c)** Vascular pO2 measured in arterioles, capillaries, and venules at each depth (0, 100, 200 µm) from 6-month-old WT and rTg4510 mice. Data are shown as mean ± SEM. Statistical analysis was performed using a mixed-effects model (REML) with mouse as a random effect and depth and genotype as fixed effects. Sphericity was not assumed, and the Geisser–Greenhouse correction was applied. Significant main effects of depth, genotype, and a depth× genotype interaction were observed in capillaries and venules only. Post hoc multiple comparisons were performed on model-predicted means with Sidak correction for multiple testing. Adjusted p-values are indicated. * p < 0.05, ** p < 0.01. **d)** 2D plane images at each depth (0, 100, 200 µm) of vascular segments imaged with 2PLIM, where point scans were used to individually measure pO2 values in different vessel types from 6-month-old WT and rTg4510 mice. **e)** Vascular pO2 measured in arterioles, capillaries, and venules at each depth (0, 100, 200 µm) from 9-month-old WT and rTg4510 mice. Data are shown as mean ± SEM. Statistical analysis was performed using a mixed-effects model (REML) with mouse as a random effect and depth and genotype as fixed effects. Sphericity was not assumed, and the Geisser–Greenhouse correction was applied. Significant main effects of depth, genotype, and a depth× genotype interaction were observed in all vessel types. Post hoc multiple comparisons were performed on model-predicted means with Sidak correction for multiple testing. Adjusted p-values are indicated. *p < 0.05, **p < 0.01, ***p < 0.01. **f)** Oxygen extraction fraction (OEF% %) in 6-month-old mice and **g)** 9-month-old mice. Violin plots show the distribution of average measurements per mouse, with the median and interquartile range overlaid. Statistical analysis was performed using a mixed-effects model (REML) with mouse as a random effect and depth and genotype as fixed effects. Sphericity was not assumed, and the Geisser–Greenhouse correction was applied. Post hoc multiple comparisons were performed on model-predicted means with Sidak correction for multiple testing. Adjusted p-values are indicated. * p < 0.05.

### Vascular pO2 is not rescued by suppressing tau production

Since the rTg4510 mouse model has repressible expression of the human tau transgene, we administered a doxycycline (DOX) diet for 3 months to our 9-month-old mice to suppress transgene expression and assess the effect on vascular oxygenation. In prior work, suppression of the tau transgene for three months reduced total soluble tau and halted neuron loss. Comparisons of the same vessels before and after DOX in a subset of mice revealed that vascular pO2 in arterioles, venules, and capillaries were unchanged after DOX administration (**Sup Fig 1A, B, C**). No effect of DOX on vessel pO2 was observed in WT mice either (**Sup Fig 1E, D, F**). Altogether, this indicates that while the presence of tau and neuron loss is associated with disturbed cerebrovascular oxygen, halting soluble tau production and neuron loss is not sufficient to impact intravascular pO2 values, at least at older ages (12-month-old).

### Tangle burden has an age-dependent relationship with vascular pO2

To assess whether intravascular pO2 values correlated with NFT burden, we used the brain-permeant fluorescent dye HS84 to label NFTs, and we calculated the mean integrated density of the HS84 signal within a 20-µm radius of vessels for which intravascular pO2 data was collected **(Fig 3A).** We then plotted the HS84 mean intensity as a function of intravascular pO2 values by vessel type, depth and age. We found a small but significant negative association between HS84 signal and intravascular pO2 in 6-month-old mice at 100 µm but not 200 µm **(Fig 3B, 3D)**. In older 9-month-old tau mice, this became a significant positive association between HS84 and pO2 values **(Fig 3C, 3E)**, suggesting that higher intravascular pO2 values correspond to vessels with greater tangle formation in the vicinity of the vessel.

**Figure 3.**
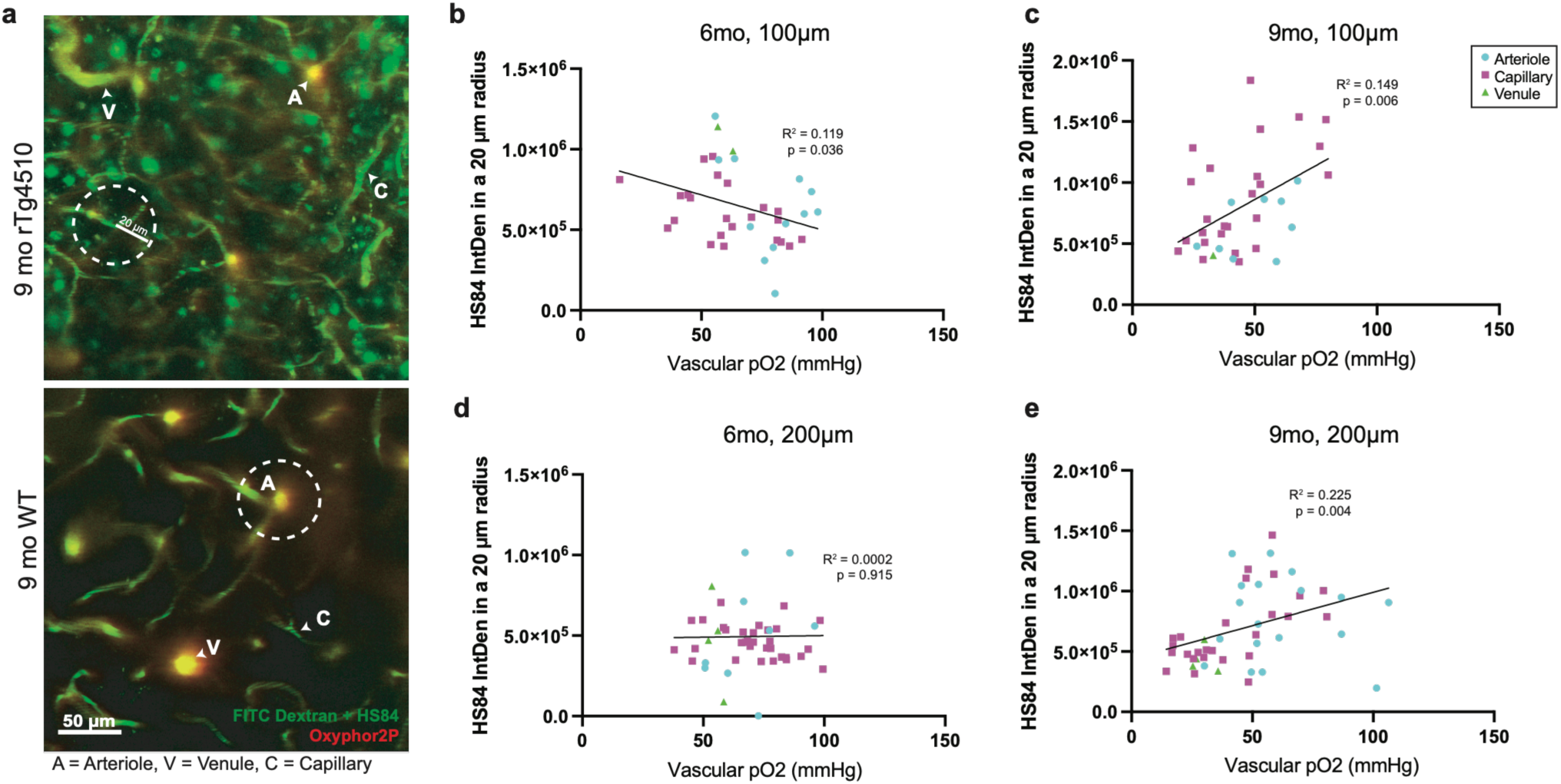
Correlation of intravascular pO2 and local tangle burden in 9-month-old rTg4510 mice. **a)** Two-dimensional images of the cortical vasculature and tangles in 9-month-old rTg4510 and WT mice at 200 µm deep into the cortex. Dotted white circles indicate the 20 µm radius area used to measure tangle density. **b)** Scatter plot of tangle density as a function of vascular pO2 (mm Hg) for 6-month-old mice at z = 100 µm. Pearson correlation analysis revealed a significant association between pO2 and tangle density (R² = 0.119, p = 0.036; n = 37 vessels). **c)** Scatter plot of tangle density as a function of vascular pO2 (mm Hg) for 9-month-old mice at z = 100 µm. Pearson correlation analysis revealed an association relationship between pO2 and tangle density (R² = 0.149, p = 0.006; n = 48 vessels). **d)** Scatter plot of tangle density as a function of vascular pO2 (mm Hg) for 6-month-old mice at z = 200 µm. Pearson correlation revealed a non-significant relationship between pO2 and tangle density (R² = 0.002, p = 0.9; n = 45 vessels). **e)** Scatter plot of tangle density as a function of vascular pO2 (mm Hg) for 9-month-old mice at z = 200 µm. Pearson correlation revealed a significant relationship between pO2 and tangle density (R² = 0.225, p = 0.004; n = 35 vessels).

Lastly, we also performed measurements in 11-month-old THY-Tau22 mice, which express soluble tau throughout the cortex and have limited NFT pathology or neuronal loss in the cortex^33^. We did not find significant differences in blood volume and hematocrit between WT and THY-Tau22 mice, but we did find a significant reduction in hematocrit in the THY-Tau22 strain compared to the rTg4510 strain **(Sup. Fig 6A, B)**. Though this line shows considerably less pathology, we observed deficits in intravascular pO2 in THY-Tau22 capillaries at every depth, including arterioles between 100-200 µm and venules at 200 µm **(Sup. Fig 4D, E, F)**, and these changes were accompanied by a trend towards increased OEF **(Sup Fig 4G)**. Collectively, these data indicate that low intravascular pO2 values are associated with the presence of tau in the mouse brain, and that these lower values are more likely to be present in areas without NFTs, particularly in aging mice.

### Tissue pO2 is decreased in 6-month-old rTg4510 mice, particularly in locations with tangles

To understand how lower intravascular pO2 values in tau-expressing mice may be impacting the availability of oxygen to cells in the brain, we performed intracranial injections of Oxyphor2P to measure cerebral tissue pO2 alongside NFTs in 6-month-old mice **(Fig 4A, 4B).** At this age, histogram plots revealed that rTg4510 mice tend to have more low pO2 values throughout the cortex, with several points measuring under 18 mmHg – values suggested by others to be near hypoxic^34,35^ (**Fig 4C)**. As seen in previous studies, lower values were observed farther from the cortical surface in both 6-month-old WT and rTg4510 mice^22^ **(Fig 4D)**. Comparisons between rTg4510 and WT mice found that overall tissue pO2 trends to be lower in tau-expressing animals at each depth, but differences between groups (WT vs rTg4510) were not significant (z = 0 µm: WT = 63.50 mmHg, rTg4510 = 44.95 mmHg, z = 100 µm: WT = 47.98 mmHg, rTg4510 = 38.29 mmHg, z = 200 µm: WT = 45.83 mmHg, rTg4510 = 33.95 mmHg) **(Fig 4D)**. Though mean tissue pO2 was not significantly different, the proportion of values that fell below 18 mmHg increased considerably with cortical depth. At z = 0 µm, 2.19% of pO2 values in WT mice are below this threshold compared to 6.84% in rTg4510 mice. Similarly, at z = 100 µm, 5.01% of WT versus 24.20% of rTg4510 pO2 values are hypoxic, and at z = 200 µm, 4.20% of WT versus 32.97% of rTg4510 pO2 values are hypoxic (p < 0.05) **(Fig 4E)**. Therefore, these results indicate that as we go deeper into the mouse cortex, the percentage of low pO2 values tends to increase and this is more extreme in rTg4510 mice.

**Figure 4.**
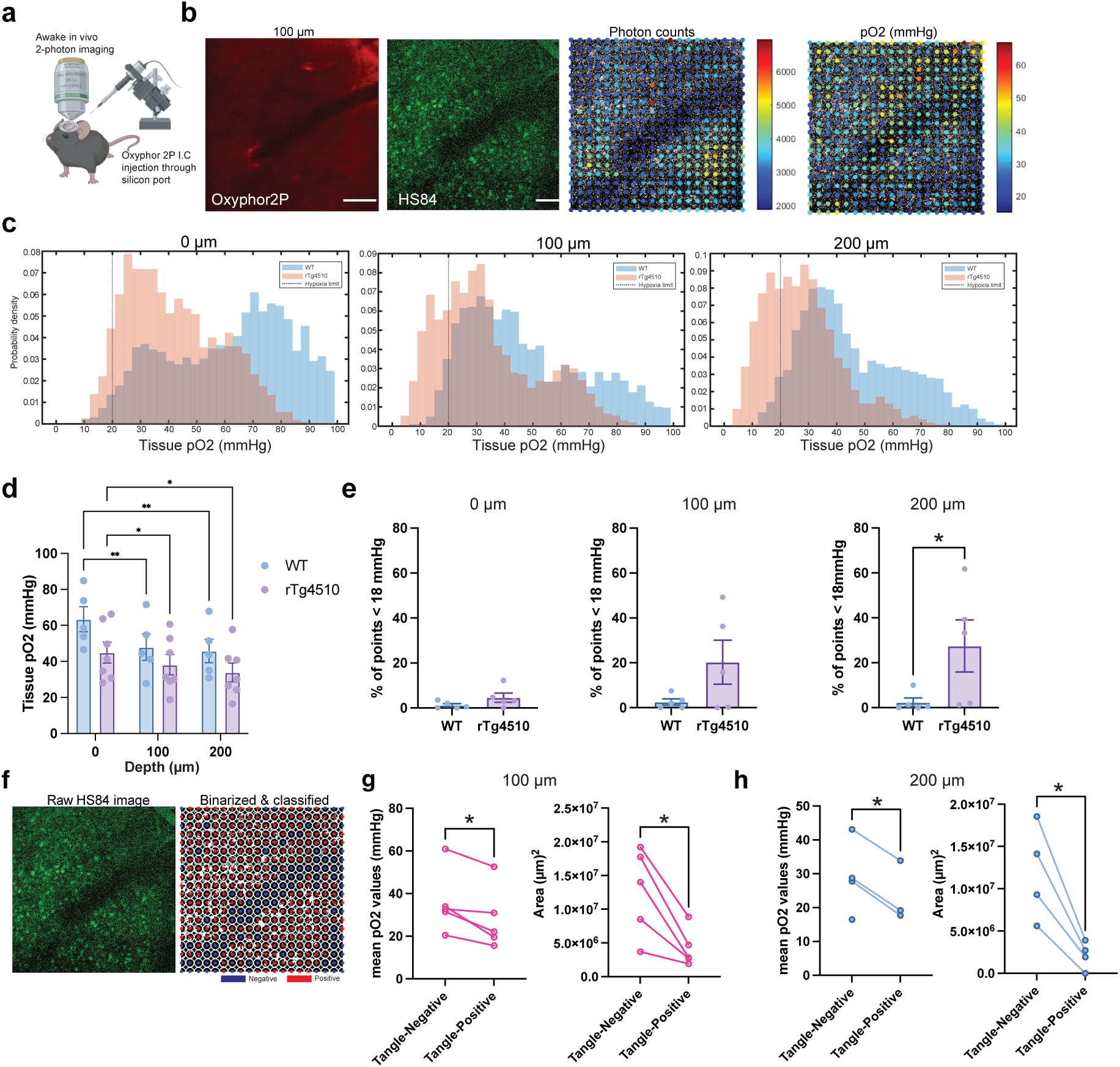
Tissue pO2 values are reduced in the presence of tau tangles in rTg4510 mice. **a)** Experimental design. Mice were injected with Oxyphor 2P through a silicon port in the cranial window and then imaged while awake using 2PLIM. **b)** Representative 2D images of rTg4510 brain cortical tissue labelled with Oxyphor2P and HS84 for tangles with its corresponding photon counts and pO2 values (mm Hg). **c)** Histograms of the distribution of pO2 values in rTg4510 and WT mice as a function of depth. Each bin set represents 400 point-scan measurements made per mouse (20 x 20 grid). A dotted black line delineates values below 18 mm Hg. **d)** Average tissue pO2 for 6-month-old WT and rTg4510 mice (n = 3 mice per group, two different sites per mouse). Statistical analysis was performed using a mixed-effect model (REML) with depth and genotype as fixed effects and subject as a random effect. Post hoc multiple comparisons of model-predicted means were performed with Sidak correction for multiple testing, and adjusted p-values of significant interactions are shown. *p<0.05. **e)** Percentage of abnormal points (pO2 values <18 mm Hg) between WT and 6-month-old rTg4510 mice at 0, 100 and 200 µm. Significance was tested using a two-paired t-test (*p<0.05). **f)** Schematic of the analysis pipeline designed to measure tissue pO2 in areas with tangles. First, raw images were binarized and thresholded to clearly visualize the signal coming from HS84 tangles. Then, the 20 x 20 pO2 grid was overlayed over the binarized image, and a circle of 13 µm radius was used as the area corresponding to the HS84+ signal. **g)** Differences in mean tissue pO2 values and area (µm^2^) between tangle “negative” and tangle “positive” regions at depth z = 100 µm and **h)** depth z = 200 µm. Each connected dot represents a measurement done in the same animal. Significance was assessed using a two-paired t-test (*p<0.05)

Prior studies have observed that neurons containing tangles are less active than non-tangle-bearing cells^36^. This feature is also observed in hypoxic neurons, and we reasoned that areas of reduced oxygen availability could exist within the cortex. To relate the distribution of tissue pO2 and to tangles, we performed a majority neighborhood classification analysis using a circular mask of 4 µm in radius to group areas positive and negative for HS84 signal around each pO2 point scan **(Fig 4F)**. We found that in tangle- positive areas, tissue pO2 is significantly lower than in tangle-negative areas at depths z = 100 µm and z = 200 µm **(Fig 4G, H)**. Altogether, this suggests that within tissue, areas with greater NFT burden exhibit greater metabolic demand, resulting in lower tissue pO2. Considering reports that rTg4510 mice exhibit neuronal silencing^36^, lower tissue pO2 may indicate that other mechanisms are driving oxygen consumption, such as increased neuroinflammation^37^, which we sought to evaluate via histology.

### Neuron loss does not occur exclusively in areas with low tissue pO2

Having found that tissue pO2 is lower in tau-expressing mice, we asked whether neurons that die are those sitting in the least oxygenated tissue. We labelled cortical neurons with AAV1-Syn-mKalama-nls, implanted a cranial window with a silicon port, and, after three weeks of recovery, injected Oxyphor2P through the port and acquired an initial tissue pO2 grid in the region where labelled neurons were located **(Fig. 5A).** Assuming that tissue pO2 remains relatively stable across multiple weeks, as seen by Hou et al^38^, the same region was then imaged weekly for four weeks, and neurons were aligned across timepoints and scored as surviving or disappearing **(Fig. 5B).** As expected, neuron loss over the four weeks was significantly higher in rTg4510 than in WT mice **(Fig. 5C).** In total, 2,282 neurons were tracked across seven animals, with 24 disappearance events: 15 of 290 neurons (5.2%) in four rTg4510 mice and 9 of 1,992 neurons (0.45%) in three WT mice, consistent with previous work^21^.

**Figure 5.**
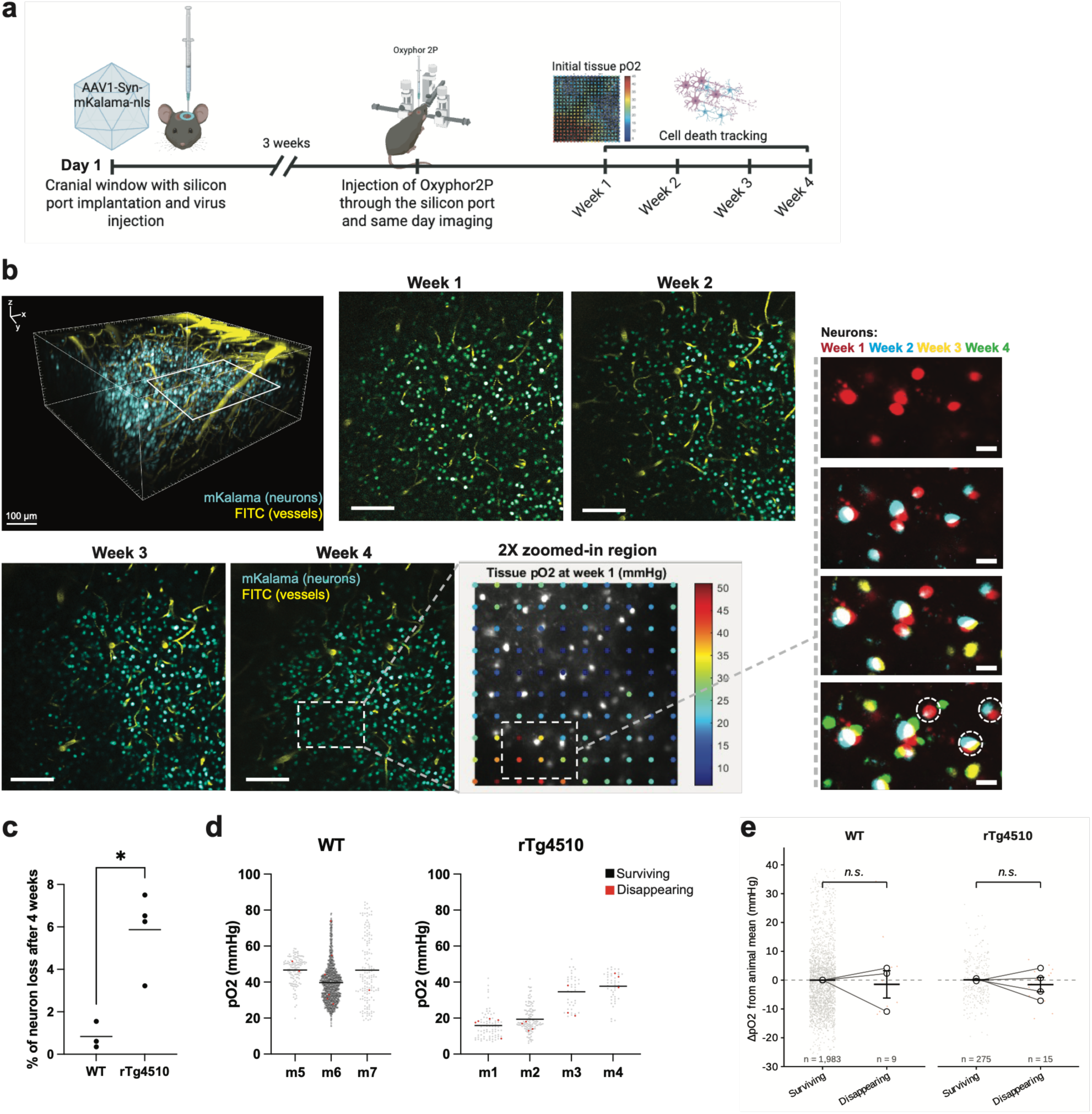
Neuron loss in mice with tissue pO2 measures. **a)** Experimental design. Mice were injected with AAV1-Syn-mKalama virus for visualizing neurons at the injection site before the implantation of a cranial window with a silicon port. On the day of imaging, Oxyphor2P was injected through the silicon port, and an initial pO2 map was acquired in the region where neurons were localized. The same region was imaged again after 1 week for four consecutive weeks**. b)** 3D rendering of neurons labeled with mKalama and blood vessels labeled with FITC. 2D images of the same region acquired over four weeks are shown, along with their corresponding pO2 maps. Neurons were tracked throughout each week as disappearing or surviving. **c)** Percentage of neuron loss per mouse in rTg4510 and WT mice (# of neurons lost / total number of neurons), lognormal Welch’s t-test (*p < 0.05). **d)** Tissue pO₂ of individual neurons in each animal, shown separately for rTg4510 (m1–m4) and wild-type (m5-m7) mice. Each point is one neuron; grey denotes neurons surviving the 4-week imaging period and red denotes neurons that disappeared. Horizontal bars indicate the per-animal mean. Neuron counts (surviving/disappearing) were: m1 75/5, m2 120/4, m3 37/3, m4 43/3; WT m5 127/2, m6 1706/6, m7 150/1; giving 2,282 neurons and 24 disappearance events across 7 animals. Baseline oxygenation differed markedly between animals (per-animal means 16.0–46.7 mm Hg), including between genotypes; surviving and disappearing neurons were therefore compared within each animal, as shown in (e). **e)** Within-animal comparison of tissue pO₂ between surviving and disappearing neurons. Each neuron is expressed as its deviation from its own animal’s mean pO₂ (ΔpO₂), removing the between-animal differences in baseline oxygenation evident in (d). Small points are individual neurons (grey, surviving; red, disappearing); open circles are per-animal means, connected within animal; black bars are the mean ± SEM across animals (rTg4510, n = 4 animals; WT, n = 3 animals). Disappearing neurons were not exclusively in low PO2 areas in either genotype (rTg4510, −1.5 ± 5.0 mm Hg; WT, −1.5 ± 8.2 mm Hg; mean ± SD across animals; within-animal permutation test, 10,000 permutations, *p* = 0.45 and *p* = 0.84, respectively). Mixed-effects logistic regression with a random intercept for animal gave odds of disappearance per 10 mm Hg of ΔpO₂ of OR = 0.77 (95% CI 0.37–1.57) in rTg4510 and OR = 1.15 (95% CI 0.69–1.94) in WT, with no evidence that the relationship differed between genotypes (interaction OR = 0.66, 95% CI 0.27–1.61, *p* = 0.36); Firth-penalized and GEE estimators gave equivalent results (interaction OR = 0.67 and 0.65). Per-animal means varied in both directions within each genotype, and with 24 disappearance events in total, these data are underpowered to exclude effects of this magnitude. n.s., not significant.

Each neuron was assigned to the pO2 value of its nearest grid point. Baseline oxygenation differed markedly between animals, with per-animal means ranging from 16.0 to 46.7 mmHg, including between genotypes **(Fig. 5D),** so surviving and disappearing neurons were compared within each animal by expressing every neuron as its deviation from its own animal’s mean (ΔpO2). Contrary to our hypothesis, disappearing neurons were not confined to tissue with low pO2 in either genotype (rTg4510, −1.5 ± 5.0 mmHg; WT, −1.5 ± 8.2 mmHg; mean ± SD across animals; within-animal permutation test, p = 0.45 and p = 0.84) **(Fig. 5E)**. Mixed-effects logistic regression gave odds of disappearance per 10 mmHg of ΔpO2 of OR = 0.77 (95% CI 0.37–1.57) in rTg4510 and OR = 1.15 (95% CI 0.69–1.94) in WT, with no evidence that the relationship differed between genotypes (interaction OR = 0.66, 95% CI 0.27–1.61, p = 0.36). Firth-penalized and GEE estimators gave equivalent results (interaction OR = 0.67 and 0.65), and per-animal means varied in both directions within each genotype **(Fig. 5E)**.

Together with our tissue pO2 measurements **(Fig. 4),** these results suggest that neuronal loss in this model parallels low tissue-level oxygen values rather than pO2 values local to neuron somas: rTg4510 mice showed both globally lower tissue pO2 and roughly tenfold higher neuronal loss than WT mice, an association that operates between animals rather than within them. With only 24 disappearance events, however, these data are underpowered to exclude a local effect, and our confidence intervals remain compatible with effects up to approximately 2.7-fold per 10 mmHg in rTg4510 mice.

### Altered tissue pO2 may be exacerbated due to an increase in astrogliosis rather than neuronal death

To better understand how in vivo measures relate to cell composition and pathological protein accumulation, we performed immunostaining of rTg4510 and WT brain tissue for neurons, tangles, microglia, astrocytes, and vasculature after imaging was concluded, and performed a layer-by-layer analysis of the same cortical region that was imaged *in vivo* **(Fig 6A)**. As expected, identification of NFTs using the phospho-tau antibody AT8 revealed a high tau burden only in rTg4510 mice **(Fig 6A).** By measuring the number of NFTs as a function of depth **(Fig 6B)**, we found that NFTs were more abundant closer to the surface of the cortex **(Fig 6C)**. By measuring the density of neurons with the pan-neuronal marker NeuN, we found that both 6- and 9-month-old rTg4510 mice have a significant reduction in neurons compared to WT **(Fig 6E, F).** We did not find any significant correlations between tissue pO2 and total neuron count **(Fig 6G)**, but tissue pO2 and AT8+ NFT counts showed a negative correlation **(Fig 6D),** indicating that areas with more tangles have less tissue pO2, which corroborates our *in vivo* results **(Fig 4E, F).**

**Figure 6.**
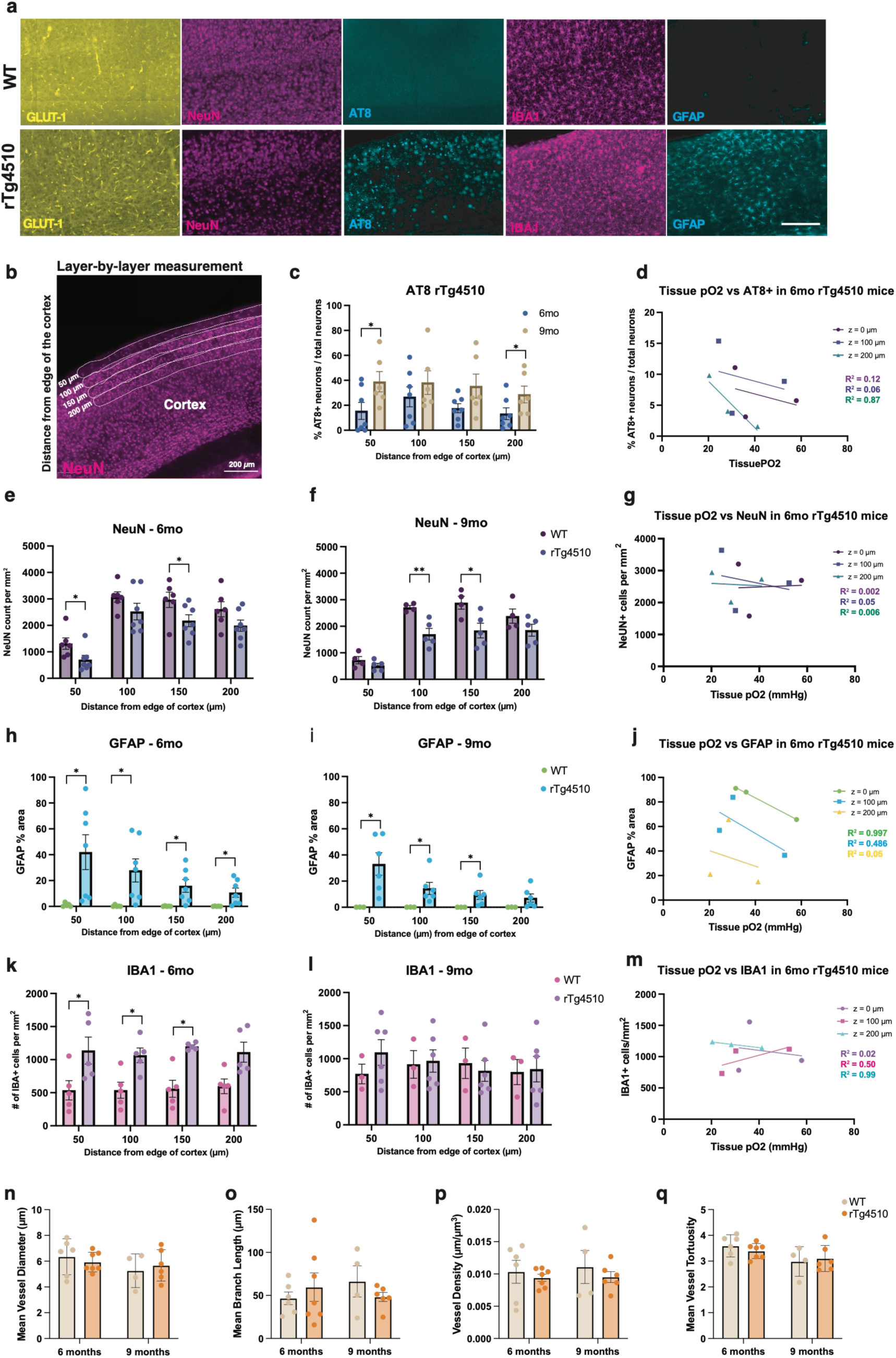
Histological correlates of altered pO2. **a)** Immunostaining of WT and rTg4510 cortex and hippocampus with NeuN, AT8, IBA-1, GFAP, and GLUT1. Scale bar = 100 µm. **b)** Example of layer-by-layer measurement performed on QuPath to quantify neurons, tangles, vessels, astrocytes, and microglia, and the distance from the edge of the cortex in µm. **c)** Percentage of AT8+ neurons/ total neurons in the brain cortex of 6-month and 9-month rTg4510 mice (n = 6-7 per group, two-tailed t-test *p<0.05). **d)** Scatter plot of tissue pO2 (mm Hg) vs AT8+ neurons/ total neurons at each depth (0, 100, 200 µm), statistical analyses were performed with Pearson correlation analysis, R^2^ and p values are shown in the graph. **e)** Quantification of neurons by depth in 6-month-old WT and rTg4510 mice (two-tailed t-test, WT n = 5, rTg4510 n = 7, *p<0.05). **f)** Quantification of neurons by depth in 9-month-old WT and rTg4510 mice (two-tailed t-test, WT n = 4, rTg4510 n = 5, *p<0.05, **p<0.01). **g)** Scatter plot of tissue pO2 (mm Hg) vs NeuN cells/mm^2^ per depth (0, 100, 200 µm). Statistical analyses were performed with Pearson correlation analysis, R^2^ and p values showed in graph. **h)** % area of GFAP in 6-month-old WT and rTg4510 as a function of depth (two-tailed t- test, *p<0.05, WT n = 5, rTg4510 n = 7). **i)** % area of GFAP in 9-month-old WT and rTg4510 as a function of depth (two-tailed t-test, *p<0.05, WT n = 3, rTg4510 n = 6). **j)** Scatter plot of tissue pO2 (mm Hg) vs % area GFAP per depth (0, 100, 200 µm). Statistical analyses were performed with Pearson correlation analysis; R^2^ and p values are shown in the graph *p<0.05. **k)** Number of IBA1+ cells/mm^2^ in 6-month-old WT and rTg4510 as a function of depth (two-tailed t-test, *p<0.05, n = 5 per group). **l)** Number of IBA1+ cells/mm^2^ in 9-month-old WT and rTg4510 as a function of depth (two-tailed t-test, *p<0.05, WT n = 3, rTg4510 n = 6). **m)** Scatter plot of tissue pO2 (mm Hg) vs IBA1+ cells/mm^2^ per depth (0, 100, 200 µm). Statistical analyses were performed using Pearson correlation analysis; R^2^ and p values are shown in the graph. *p<0.05. **n)** Mean vessel diameter, **o)** Mean branch length, **p)** Vessel density, **q)** Mean vessel tortuosity for WT and rTg4510 mice at 6 and 9 months old. No significant differences were seen between groups (two-tailed t-tests, *p<0.05, n = 5-6 per group).

We also found that rTg4510 mice show a significant increase in reactive astrocytes (GFAP % area) in brain cortex compared to WT at both 6 and 9 months old **(Fig 6H, I)** and a significant increase in microglia (IBA1 positive cells) in 6-month-old mice but not 9-month-old mice **(Fig 6K,L).** We did not find any association between IBA1 and tissue pO2 in rTg4510 mice **(Fig 6M),** but GFAP % area showed a consistent negative association with tissue pO₂ across all depths; however, these relationships did not reach statistical significance **(Fig 6J).** The association between reactive astrogliosis and lower tissue pO₂ may reflect underlying neurovascular dysfunction associated with disease progression.

We did not find significant differences in vascular metrics measured with GLUT-1 staining, such as vessel diameter, tortuosity, branch length or density **(Fig 6N, O, P, Q)**. At the ages we explored (6 and 9 months), vascular architecture remains similar to WT mice; however, at older ages (10+ months), changes in vascular architecture, such as vessel tortuosity, diameter, and staling, become more significant, as seen in previous studies^39^. We do not rule out the possibility that what we observe in vascular architecture *ex vivo* is also present *in vivo*, particularly given that differences in blood flow and vessel dilation between WT and rTg4510 mice may arise at some point during their lives.

We also conducted immunostaining of brain tissue in 11-mo THY-Tau22 and WT mice **(Sup Fig. 5A)**. At this age, few NFTs were present in THY-Tau22 mice, with the majority appearing in deeper cortical layers, as expected **(Sup Fig 5B, C).** Unlike in the rTg4510 mouse line, in THY-Tau22 mice neuronal loss is not readily apparent compared to WT **(Sup Fig 5D, E).** Similarly, THY-Tau22 and WT mice had similar densities of microglia and reactive astrocytes **(Sup Fig 5G, I)**, with only a slight elevation in microglial number detected at 200 µm **(Sup Fig 5H).** Altogether, the limited changes in cell composition and the lack of NFTs in these mice suggests that these factors are unlikely to be contributing to observed differences in intravascular pO2.

## DISCUSSION

In this work, we set out to better understand the connection between tau pathology and vascular dysfunction in disease by directly measuring intravascular and tissue oxygen using two photon imaging of awake mice. We found that tau pathology is associated with significant alterations in brain oxygenation. Disrupted pO2 was evident in two separate mouse models that develop different extents of tau pathology and neuron loss. While low pO2 values were common in tau mice, particularly in areas with greater NFT burden, they did not fully explain the loss of individual neurons. Together, these findings are important for determining how vascular and metabolic dysfunction occur alongside AD and contribute to neurodegeneration.

The impact of tau pathology and brain oxygenation was most evident in 6-month-old mice, where we collected both intravascular and tissue pO2 measures alongside tracking tangles and neurons. At this age, overall intravascular pO2 measures were elevated or equivalent (depending on cortical depth) in rTg4510 mice compared to wild type, but locations with greater burden of NFTs showed reductions in both intravascular and tissue pO2 values. This relationship suggests that the development of tau pathology is closely linked to altered oxygen metabolism within the surrounding tissue. Importantly, pO2 was not correlated with histological measures of neuron density. Instead, lower oxygen levels may reflect increased metabolic demand from the gliosis accompanying tau pathology. In line with this possibility, mice at the ages examined also had increased GFAP+ astrocytes and GFAP area was negatively correlated with tissue pO2 values. A previous study by Liu et al.^37^, which used mice that developed amyloid plaques (APP/PS1), similarly found that neuroinflammation increases oxygen extraction fraction, and others have reported that glia contribute significantly to brain energy expenditure^40^. Thus, the reduction in pO2 observed in regions with abundant tau pathology may be driven by activated glial cells rather than simply reduced oxygen delivery or changes in neuronal metabolic activity.

Despite this association between tau pathology and tissue oxygenation, our cell-level analysis did not support a simple model in which local low pO2 values directly cause the death of individual neurons. When we longitudinally tracked 2,282 neurons, neurons that subsequently disappeared were not preferentially located in areas with lower tissue pO2. With only 24 disappearing neurons observed in this cohort, while we cannot rule out that a cell-level relationship does not exist, the absence of a relationship was observed across four different approaches to estimating the local oxygen environment. Our design also assumes that tissue pO2 remains stable over weeks^38^, and assigns each neuron the value of its nearest grid point, so slow drifts in oxygenation or an acute decline preceding death would go undetected. In sum, although tau mice with notably lower tissue pO2 exhibited greater neuron loss compared to wild-type controls, areas of low pO2 do not appear to predict which individual neurons are lost in following weeks.

These findings suggest that abnormal oxygenation and neurodegeneration are linked at the level of the tissue rather than at the level of individual neurons. One possibility is that low pO2 associated with tau pathology reflects increased oxygen consumption by reactive glia and other activated cell types, while neuron loss represents a related but distinct consequence of disease progression. This interpretation is consistent with the observation that oxygen diffuses over tens of microns^41^, such that the oxygen environment experienced by an individual neuron is determined by its broader capillary neighborhood rather than by its immediate surroundings. It would therefore be possible for neurons exposed to a similar oxygen environment to have different fates. In line with this idea that neurons are supported by relatively large territories, investigators who experimentally occluded blood vessels showed that a single capillary occlusion does not induce damage to a nearby neuron, and multiple capillary occlusions were necessary to trigger cell apoptosis^42^. Last, we note that in this work we considered a threshold of 18 mmHg as regions of low tissue pO2 based on prior reports that investigated hypoxia, but the amount and duration of reduced oxygenation required to contribute to neuron death remains uncertain, particularly in disease states, and additional studies would be required to better define the conditions that drive cell fate under low oxygen.

Importantly, we also investigated whether these changes in oxygen metabolism were reversible. However, even after a 3-month diet with doxycycline to shut off tau transgene expression^43^, vessel pO2 remained unchanged from pre-treatment levels. By contrast, previous work shows that tau suppression recovers other disease-associated consequences such as reduced neuronal activity^36^, microangiopathy^44^ and cerebral blood flow^45^. Age differences likely contribute to our observed lack of reversal since we began treatment in 9-month-old mice; at this age, mice exhibited greater neuron loss, tau pathology and altered relationships between intravascular pO2 and NFTs than what was seen in 6-month-old animals. This may indicate that the effects of tau on delivery of oxygen to the brain may become too advanced, prohibiting rescue by reducing tau burden alone. These findings are also consistent with our prior work showing upregulation of vascular inflammation markers and altered vessel structure are also not easily reversed in these later stages in these mice^44^.

Contrary to our expectations, we observed even more pronounced changes in vascular oxygenation in 11-month-old THY-Tau22 mice. This was surprising given that the model exhibits considerably less pathology than the rTg4510 line, even at this older age. At every depth, capillary pO2 was significantly lower than WT mice and arteriolar pO2 was lower at z = 100 µm, 200 µm. While blood volume was similar, we found that THY-Tau22 mice have a lower blood hematocrit than rTg4510 mice. Age could also be a significant factor influencing this, compromising oxygen delivery to the brain from the lungs^46^, as seen in humans^47^. This reduced oxygen carrying capacity could be one explanation for the reduced intravascular pO2 measures in these mice, considering arteriolar measures were also impacted. Altogether, different factors may drive reduced pO2 in each of these tau mouse models, but abnormal intravascular pO2 was readily observed in both.

A limitation of this study is that we only measured pO2 in awake mice at rest. Awake imaging also allowed us to perform measurements without the confounding effects of rodent anesthesia on cerebral metabolism and blood flow^48^. In future studies, it would be interesting to measure the changes in oxygenation during a functional task, especially since AD mice have been shown to have impaired responses to visual and limb stimulation^25^. Such impairment in functional hyperemia may exacerbate any oxygen deficits and neuronal death. Pairing this data with blood flow measures in all vessel types can offer a deeper understanding of the mechanisms driving abnormal vascular and tissue oxygenation, such as altered capillary transit time heterogeneity^32,49^ and cerebral vascular reactivity^50^.

In all, our findings indicate that brain oxygen metabolism is strongly influenced by NFT accumulation and associated gliosis, emphasizing the complex relationship between pathological protein accumulation, cellular metabolism, and neurodegeneration. In human neuroimaging studies, the cerebral metabolic rate of oxygen and OEF can be measured noninvasively, although reported changes in AD have been mixed^51–56^, potentially reflecting dynamic relationship observed here. These findings also support incorporating measures of oxygen metabolism into therapeutic outcome assessments, as vascular dysfunction may persist even when the initiating pathology is reduced. Together, by providing essential cell-level physiological data, these results highlight brain oxygen metabolism as a component of the tissue environment that shapes neuronal vulnerability in AD.

## ACKNOWLEDGMENTS

We thank Marcel Weinreich for support with the Crowntracker software for awake imaging.

## AUTHOR CONTRIBUTIONS

These experiments were designed and conducted by NERU and RB. NERU performed cranial window surgeries, Oxyphor2P and mKalama injections, and imaging. MT and MA provided code to quantify phosphorescence decays. NW, DS and SEM performed immunostaining, image analysis and quantification. AD habituated mice to awake imaging. EG performed ilastik segmentation and quantification of blood vessel metrics. TZ wrote code for neuronal death alignment with tissue pO2 and statistical analysis in MATLAB. SS offered guidance and support with the microscope and oxygen data. SB synthesized the phosphorescent probe Oxyphor2P. SV performed calibrations of Oxyphor2P to enable quantitative pO2 measurements. NERU prepared figures for the manuscript and wrote the paper with RB. All authors reviewed the paper.

