## Supplementary material for "Depth-Dependent Alterations in Cerebral Blood and Tissue Oxygenation Associated with Tau in Alzheimer’s Disease Models"

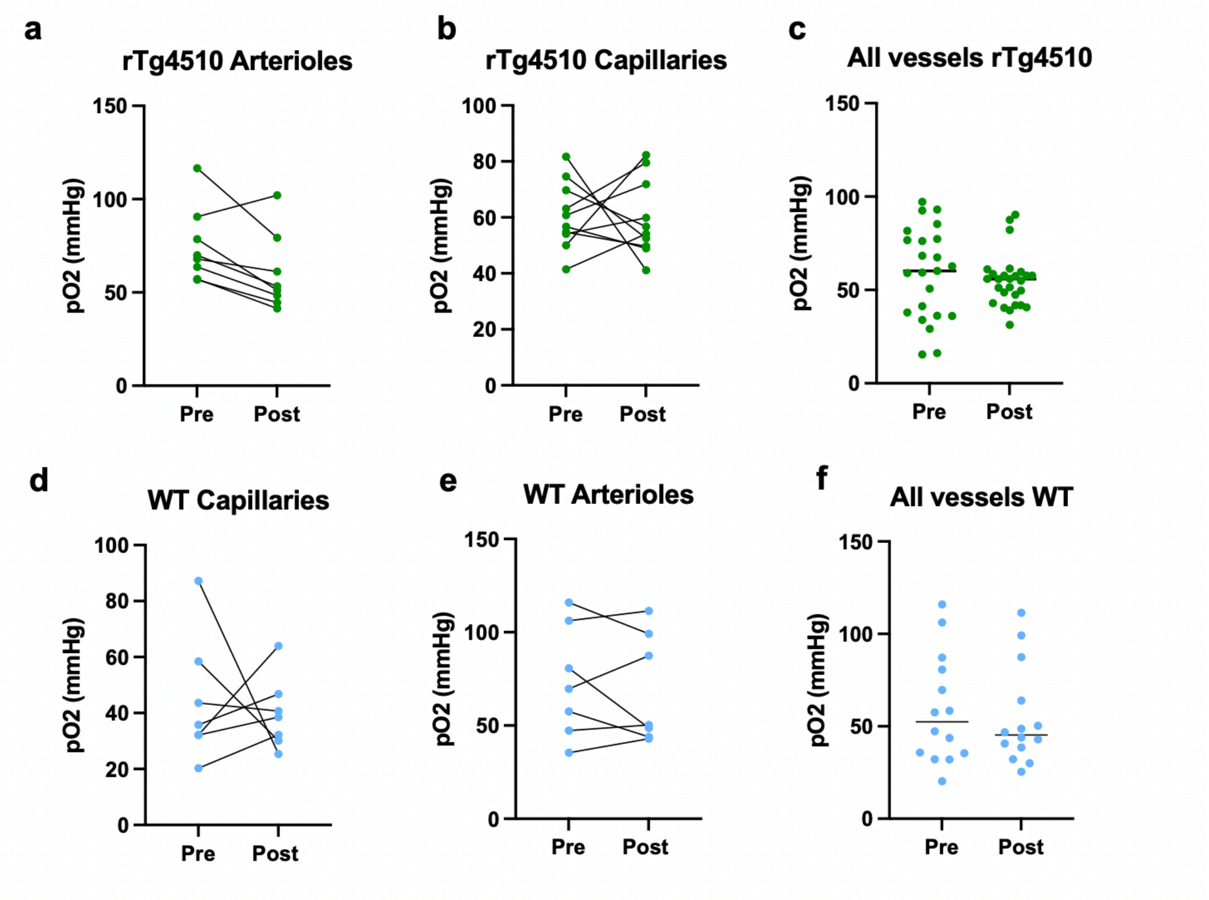


**Supplementary Figure 1.** Doxycycline (DOX) does not affect vascular pO2 values in 12-month-old rTg4510 and WT mice. **A)** pO2 values in arterioles, **B)** capillaries and **C)** all vessels in rTg4510 mice pre and post DOX treatment. 9-month-old mice were fed a DOX diet at the end of the first imaging session, and vascular pO2 was then measured after 3 months on the same blood vessels. **D)** pO2 values in capillaries, **E)** arterioles and **F)** all vessels in WT mice before and after doxycycline treatment. Each dot represents one vessel, and the same vessel was measured before and after treatment. Black lines connecting dots indicate measurements performed in the same vessels.

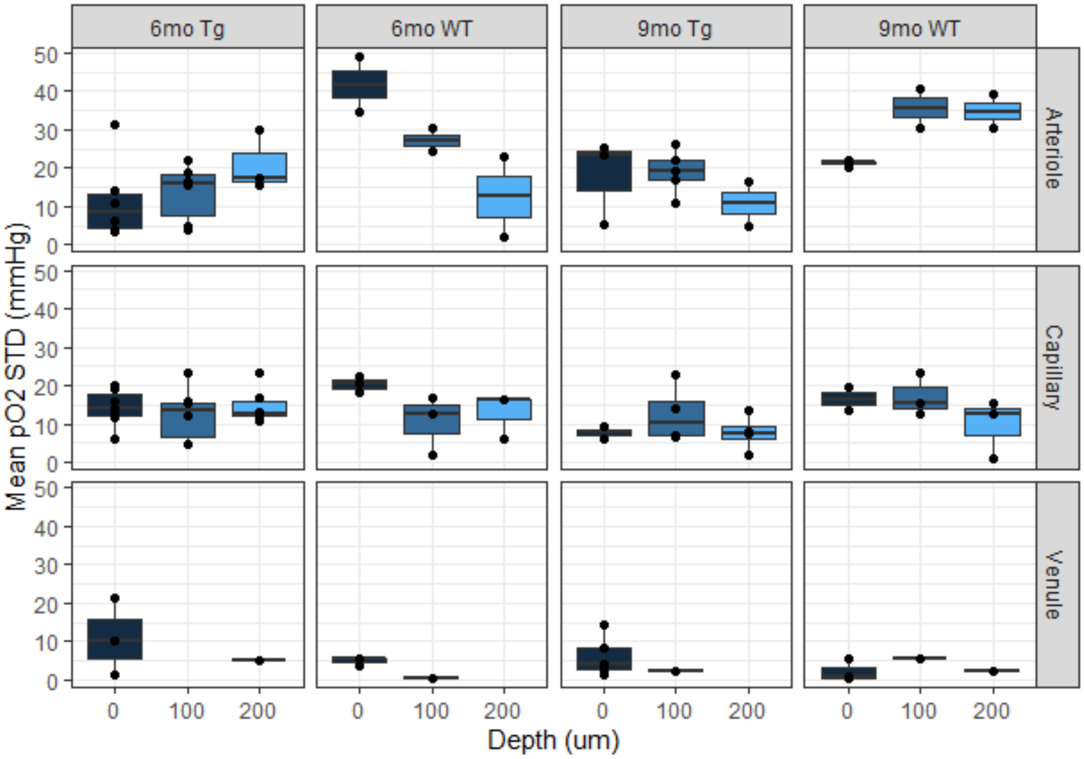


**Supplementary Figure 2.** Standard deviations of the mean pO2 values in arterioles, venules and capillaries of 6- and 9-month-old WT and rTg4510 mice.

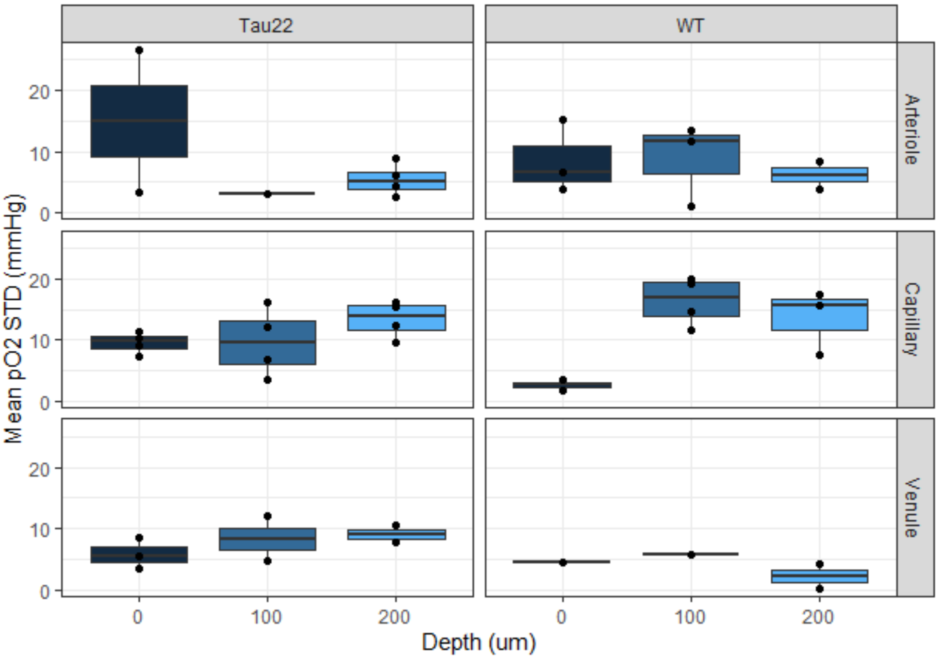

**Supplementary Figure 3.** Standard deviations of the mean pO2 values in arterioles, venules and capillaries of 11-month-old WT and THY-Tau22 mice.

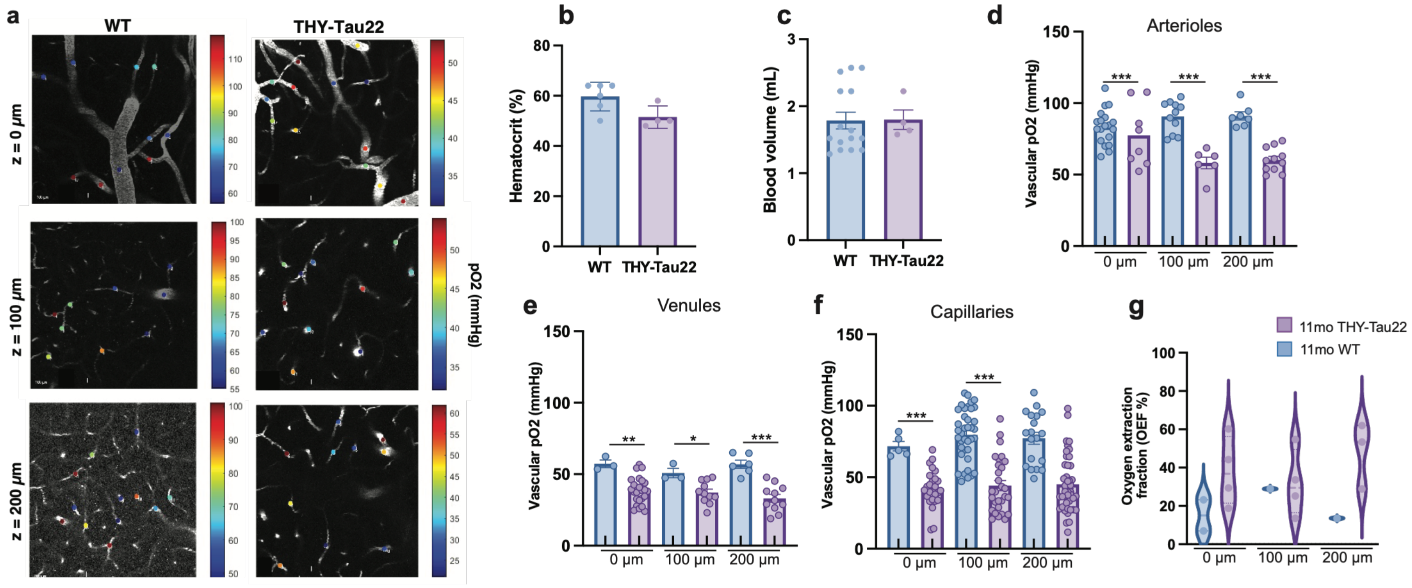


**Supplementary Figure 4. A)** 2D plane images at each depth (0, 100, 200 µm) of vascular segments imaged with 2PLIM where point scans were used to individually measure pO2 values in different vessel types from 11-month-old WT and THY-Tau22 mice. **B)** Hematocrit and **C)** blood volume from Tg4510, THY-Tau22 and WT mice. **D)** Vascular pO2 in arterioles, **E)** venules and **F)** capillaries in WT and THY-Tau22 mice at 0, 100, 200 µm. Each point represents a vessel. Significance was achieved via two-way ANOVA followed by Tukey's multiple comparison test. Capillaries (WT vs THY-Tau22 z=0: p = 0.0096, z=100: p = 0.008, z=200: p = 0.0046), Arterioles (WT vs THY-Tau22 z=0: p = 0.2739, z=100: p = 0.0002, z=200: p = 0.0069). Venules (WT vs THY-Tau22 z=0: p = 0.06, z=100: p = 0.18, z=200: p = 0.0032). **G)** Oxygen extraction fraction of 11mo WT and THY-Tau22 mice at 0 µm, 100 µm, and 200 µm. Significance was tested via two-way ANOVA.

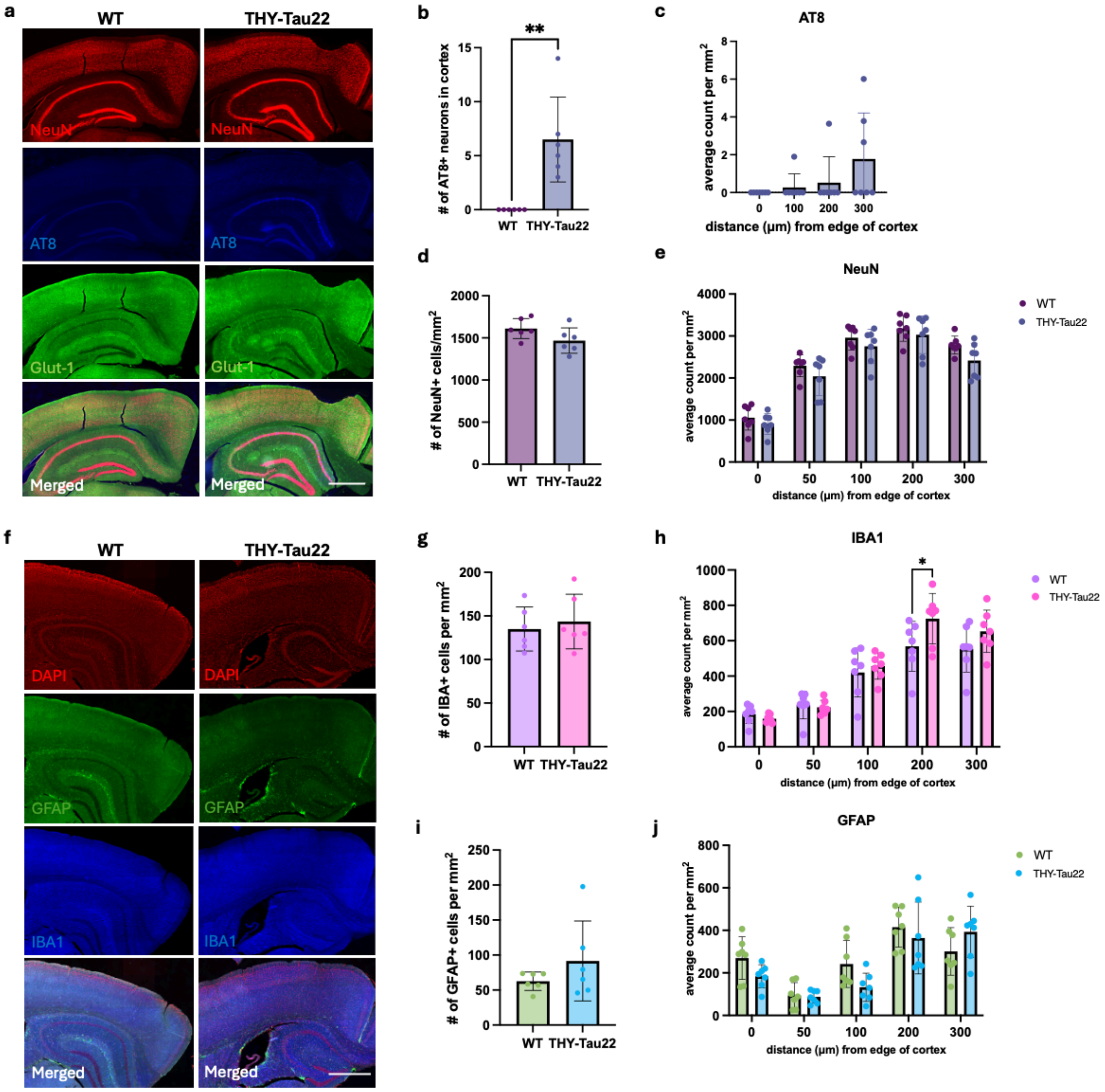


**Supplementary Figure 5**. **A)** Immunostaining of WT and THY-Tau22 cortex and hippocampus with NeuN, AT8 and Glut-1. Scale bar = 1 mm. **B)** Quantification of the number AT8+ neurons in brain cortex of WT and THY-Tau22 mice (* $p < 0.05$, **$p<0.01$, two-tailed t-test, n=5-6). **C)** Quantification by depth of AT8+ cells per $mm^2$ in THY-Tau22 mice (n = 5-6), **D)** Total number of neurons in WT and THY-Tau22 mice (two-tailed t-test, n = 5-6). **E)** Quantification of neurons by depth in WT and THY-Tau22 mice (two-way ANOVA, depth: $F (4, 48) = 204.1$, $P<0.0001$, phenotype: $F (12, 48) = 8.406$, $P<0.0001$, post-hoc comparisons non-significant). **F)** Immunostaining of WT and THY-Tau22 cortex and hippocampus with DAPI, GFAP and IBA1. Scale bar = 1 mm. **G)** Total number of IBA1+ cells/$mm^2$ in WT and THY-Tau22 (*$p < 0.05$, two-tailed t-test, n = 5-6). **H)** Number of IBA1+ cells/$mm^2$ in WT and THY-Tau22 as a function of depth (mixed effect models, depth: $F (4, 48) = 124.7$ $P<0.0001$, phenotype: $F (12, 48) = 6.460$, $P<0.0001$, Sidak multiple comparison test: WT vs THY-Tau22 at 200 um $p = 0.0322$, n = 5-6). **I)** Total number of GFAP+ cells/$mm^2$ in WT and THY-Tau22 (two-tailed t-test, n = 5-6). J) Number of GFAP+ cells/$mm^2$ in WT and THY-Tau22 as a function of depth (mixed effect models, F (4, 48) = 29.29, P<0.0001, phenotype: F (12, 48) = 3.032, P=0.0031, Sidak multiple comparison test shows no significant differences).

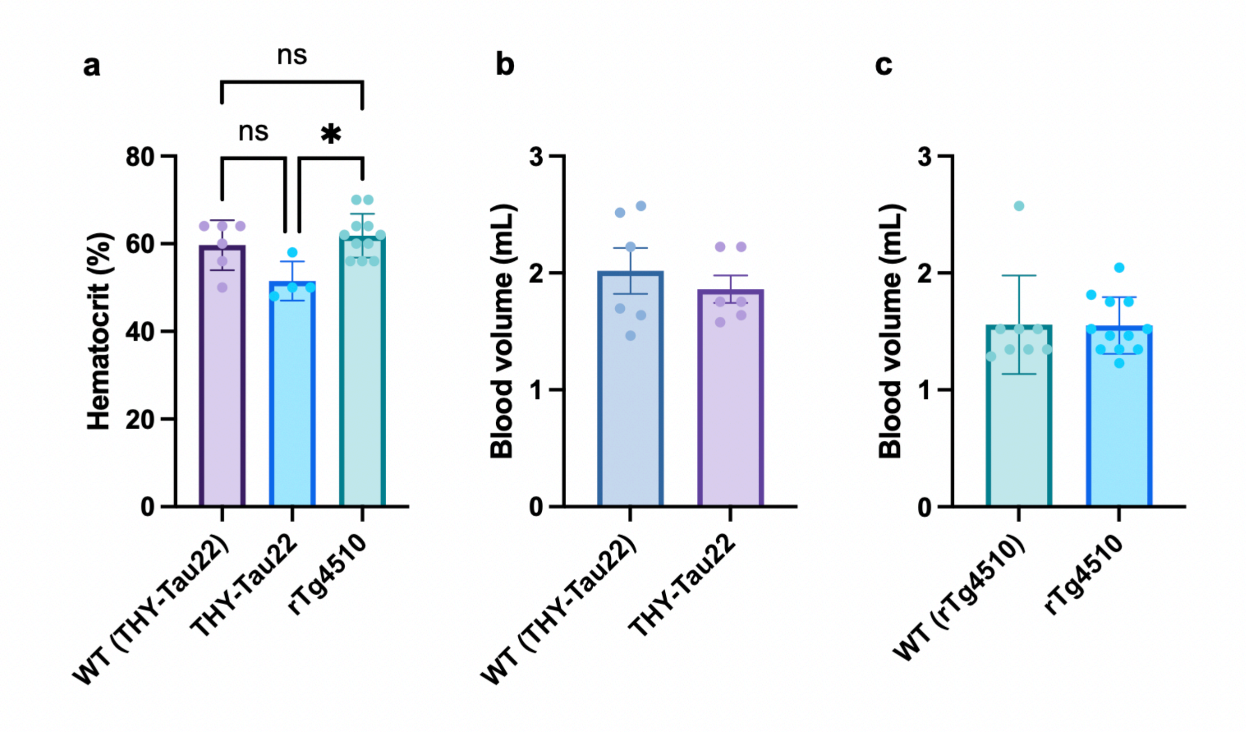


**Supplementary Figure 6**. **A)** Hematocrit from all groups combined, **B)** blood volume from WT and THY-Tau22 and **C)** blood volume from rTg4510 and WT mice. *p <0.05, Kruskal-Wallis test.
